# Memory T and B cells with recognition of avian influenza hemagglutinins are poorly responsive to existing seasonal influenza vaccines

**DOI:** 10.1101/2025.04.28.651131

**Authors:** Christopher A Gonelli, Marios Koutsakos, Robyn Esterbauer, Ming Z M Zheng, Yee-Chen Liu, Amanda Zin, Lara S U Schwab, Danielle Tilmanis, Malet Aban, Aeron Hurt, Stephen J Kent, Jennifer A Juno, Adam K Wheatley

## Abstract

Immunisation remains the most cost-effective mechanism to combat global influenza infection and is widely employed against seasonal influenza viruses. Zoonotic transmission of avian influenza A viruses represents a significant threat to human health given the lack of population level immunity, which could translate into an influenza pandemic. Therefore, there is a need to better understand pre-existing human immunity against avian influenza strains. as highlighted by the recent rapid, global spread of avian H5Nx clade 2.3.4.4b variants. Here, we sought to quantify the frequencies and specificities of B cells recognising avian hemagglutinin (HA) within unexposed adults, and to characterise the ability of seasonal immunisation to boost cross-reactive immune responses to H5Nx strains, including from clade 2.3.4.4b. Low but detectable serum antibody titres against H5 and H7 avian influenza HA were observed in donors. The frequency of memory B cells with cross-reactive recognition of H5 and H7 HA was low and 2–5 fold lower than populations of seasonal H1N1 and H3N2 HA-specific B cells. Boosting of B cell responses against H5Nx clade 2.3.4.4b HA following seasonal immunisation were sporadic with only 3 out of 19 individuals showing an increased population of probe-positive cells. Cross-reactive B cells generally expressed immunoglobulins drawn from variable heavy chain genes associated with recognition of the HA stem (VH6-1, VH1-69, VH1-18). CD4+ T cell responses towards H5 HA were also weakly boosted with little to no increase in circulating T follicular helper cell populations. These findings highlight the need for avian influenza-specific vaccine products to bolster immunity in human populations, with consideration for use in pre-pandemic preparedness to expand baseline frequencies of avian influenza-specific memory B and T lymphocytes.

## Introduction

Zoonotic transmission of avian influenza A viruses such as H5N1 and H7N9 is often associated with extremely high pathogenicity and case fatality rates in humans^1–5^. Due to a lack of population level immunity, cross-over from avian reservoirs represents a pressing and emergent threat to human health, with any global pandemic likely to be associated with large loss of life and significant social upheaval. While a variety of monovalent vaccines targeting H5N1 or H7N9 influenza have been produced and are immunogenic in humans^6–10^, the unpredictable location and timing of any emergent pandemic makes antigenic mismatch between existing vaccines and/or vaccine seed stocks highly likely. There is therefore a need to better understand pre-existing human immunity against avian influenza strains, particularly in light of the recent global spread of avian H5Nx 2.3.4.4b variants^11^.

The existence within unexposed individuals of serum antibodies able to bind and/or neutralise the hemagglutinin (HA) of avian influenza strains has been widely reported. H5- and/or H7-reactivity is observed in both pooled intravenous immunoglobulin (IVIG) preparations^12,13^ and in serum samples from human cohorts^14–16^, although reported serological concentrations are generally very low. In addition, monoclonal antibodies (mAbs) binding H5 or H7 isolates are readily isolated from subjects not directly exposed to avian influenza by immunisation or infection^17–19^ and can protect against H5N1 or H7N9 challenge in mice^17,18,20^. Notably, immune exposure to antigenically divergent HA drives the preferential expansion of highly cross-reactive antibody and memory B cell populations^21–24^, including those expressing rare mAbs able to neutralise and/or protect against both group 1 and group 2 influenza^25,26^. Thus, the human immune system appears highly capable of targeting conserved epitopes shared by seasonal and avian influenza strains.

Existing H5-specific cellular responses are also present within populations of unexposed individuals, as evidenced by CD4+ (and CD8+) memory T cell reactivity predominantly recognising internal proteins of avian viruses^27–31^. This cross-reactive memory is likely the product of epitope conservation between seasonal and avian viruses^32^, and can be expanded following inactivated H5N1 virus immunisation ^33^. Studies suggest that the immunogenicity of HA-based vaccines in humans is determined, in part, by levels of pre-existing HA-specific CD4 memory T cells^34,35^. In populations with low baseline H5- or H7-reactive T cell pools, vaccine immunogenicity may be improved by prior CD4 T cell priming^33^ or by covalent coupling of novel HA antigens to seasonal HA proteins^36^.

Most recently novel H5 viruses from clade 2.3.4.4.b have spread globally through wild bird populations in four continents^37^, and with zoonotic outbreaks have reported within domestic cattle^38–40^, sea mammals^41,42^ and mustelids^43^. Disease course within mammals varies in severity, with a mild disease largely confined to mammary tissues reported in cattle, while infection in cats^44^ and experimentally infected naïve ferrets^45^ and non-human primates^46^ can be highly pathogenic causing major lung pathology and/or death. To date, zoonotic infections in humans have been nearly universally mild, likely reflecting a degree of cross-protective immunity within the population seeded by seasonal exposure.

Here we sought to quantify the frequencies and specificities of B cells recognising avian HA within adults, and to determine the extent to which seasonal immunisation can boost cross-reactive immune responses to H5 including lineages such as 2.3.4.4b. We find memory B cells recognising HA from avian H5N1 and H7N9 influenza strains are widely prevalent in healthy unexposed Australian adults and utilise stereotypic immunoglobulin sequences previously shown to be able to protect in animal models. Vaccination with seasonal inactivated influenza vaccine drives a modest, transient expansion of B cells and CD4+ T cells recognising avian influenza strains. Our results suggest that while population level immunity to H5N1 2.3.4.4b in the form of antibody and memory lymphocyte populations is widespread, targeted vaccine strategies against H5N1 will be required to markedly bolster immunity to emerging avian influenza threats.

## Results

### Antibodies and B cells binding hemagglutinin from H5 and H7 avian influenza strains are widely prevalent in unexposed adults

Consistent with previous reports^14–16^, we observed low but detectable serum antibody titres reactive against H5 and H7 avian influenza strains within a cohort of healthy, unexposed Australian adults (N=18), a level 5- to 118-fold reduced compared with endemic H3N2 and H1N1 influenza strains (Fig. 1A). Memory B cells recognising HA from historical H5 (H5N1; A/Indonesia/05/2005) and H7 (H7N9; A/Shanghai/02/2013) avian influenza strains were quantified by flow cytometry using recombinant HA probes^47^. Distinct H5+ and H7+ memory B cell populations were detected in all individuals tested (Fig. 1B), with median respective frequencies of 0.054% (range 0.021 - 0.122) and 0.056% (range 0.031 - 0.086) of total class-switched (IgD-) B cells (Fig. 1C). Cross-reactive memory B cells binding both H5 and H7 probes were comparatively infrequent (0.008%; range 0.002 - 0.046), however notable populations were readily discernible in several subjects screened (Fig 1B-C). As a point of reference, memory B cells specific for seasonally epidemic H1N1 (A/New Caledonia/20/1999 and A/California/04/2009) and H3N2 (A/Hong Kong/1/1968 and A/Victoria/361/2011) influenza strains were commonly observed at frequencies 2–5 times greater within the same individuals (Fig. 1D) (N=16 matched subjects). Memory B cells specific for avian HA were phenotypically comparable to the parental memory B cell population, displaying a similar distribution of surface immunoglobulin usage (Supplementary Fig. 1A) and a predominantly resting memory phenotype (CD27+ CD21-) (Supplementary Fig. 1B). There was a weak association between subject age and the frequency of cross-reactive memory B cells, an observation that requires clarification in larger cohorts (Supplementary Fig. 1C).

**Figure 1:**
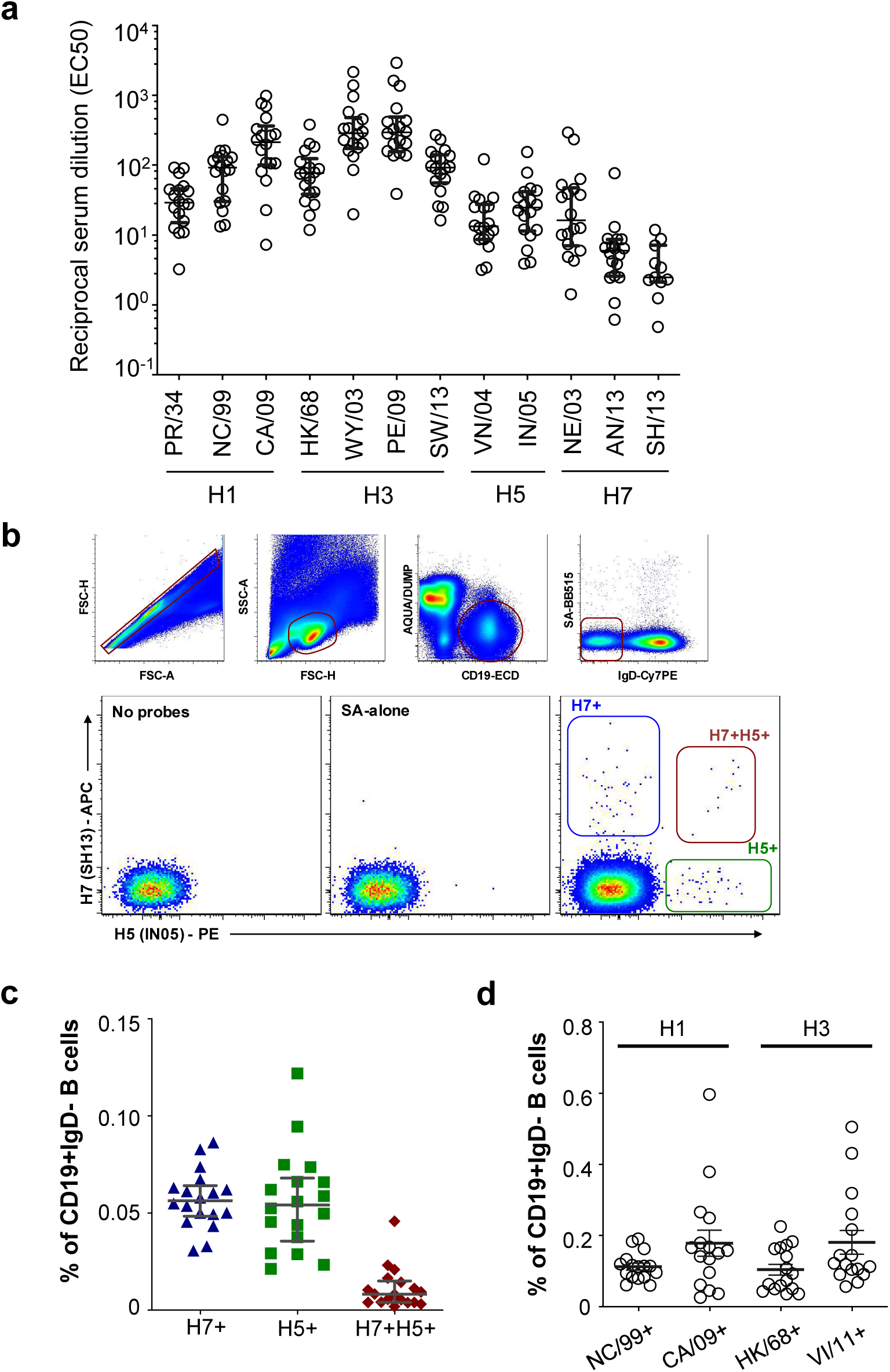
Serum antibody and memory B cells recognising avian influenza strains are widely prevalent. (**A**) Plasma samples from healthy volunteers (N=18) were screened by ELISA for reactivity against HA from diverse influenza A strains. Reciprocal serum dilutions yielding half maximal binding (EC50) for each antigen are shown. **(B)** Staining of cryopreserved PBMCs allows identification of CD19+ IgD- B cells not binding a streptavidin-BB515 decoy. Co-staining with recombinant HA probes derived from H7N9 A/Shanghai/01/2013 (SH13) and H5N1 A/Indonesia/5/2005 (IN05) delineates single- and cross-reactive memory B cell populations. (**C**) Frequencies of H5+, H7+ or H7+H5+ memory B cells in healthy volunteers (N=18). (**D**) Frequencies of memory B cells binding seasonal H1N1 (NC99 and CA09) and H3N2 (HK68 and VI11) influenza strains were measured in healthy volunteers (N=16). Lines indicate median and IQR. PR/34, A/Puerto Rico/8/1934; NC/99, A/New Caledonia/20/1999; CA/09, A/California/04/2009; HK/68, A/Hong Kong/1/1968; WY/03, A/Wyoming/3/2003; PE/09, A/Perth/16/2009; SW/13, A/Switzerland/9715293/2013; VN/04, A/Vietnam/1203/2004; IN/05, A/Indonesia/5/2005; NE/03, A/Netherlands/219/2003; AN/13, A/Anhui/01/2013; SH/13, A/Shanghai/01/2013; VI/II, A/Victoria/361/2011

### Recovered immunoglobulins recognise HA from diverse influenza subtypes

Single memory B cells binding H5, H7 or both HA probes were sorted from two individuals and immunoglobulin genes sequences recovered as previously described^47,48^. A small panel of monoclonal antibodies was expressed (Fig. 2A) and binding to HA from diverse influenza strains was confirmed by ELISA using recombinant HA proteins (Fig. 2B). Consistent with reports of broadly cross-reactive human antibodies, mAbs derived from H5 specific B cells primarily bound influenza A viruses from Group 1, while those from H7-specific B cells bound group 2. Antibodies from B cells with H5/H7 cross-binding activity bound more broadly and were generally drawn from IGHV6-1 and IGHV1-18 convergent classes previously described^26,49^. The ability of the mAbs to neutralise virus activity was assessed using hemagglutination inhibition (HAI) and focus reduction assays (FRA) against a panel of influenza A and B viruses (Fig. 2C). None of the isolated mAbs showed HAI activity against H1, H5, H3 or H7 viruses nor influenza B viruses. However, most of the mAbs showed FRA activity against one or more HA subtypes, particularly for H5 among the H5/H7 cross-reactive mAbs. As HAI exclusively measures antibodies targeting the HA head domain while FRA activity measures antibodies that inhibit virus spread (binding, fusion and release), this suggests that as expected mAbs are specific for the stem region of HA. This is consistent with the positive control stem-binding mAb, CR9114, showing no HAI activity while having broad FRA activity against the influenza A viruses^50^. Overall, pre-existing neutralising antibodies binding the HA stem domain are prevalent among unexposed individuals and can exhibit neutralising and potentially protective activity against avian influenza strains.

**Figure 2.**
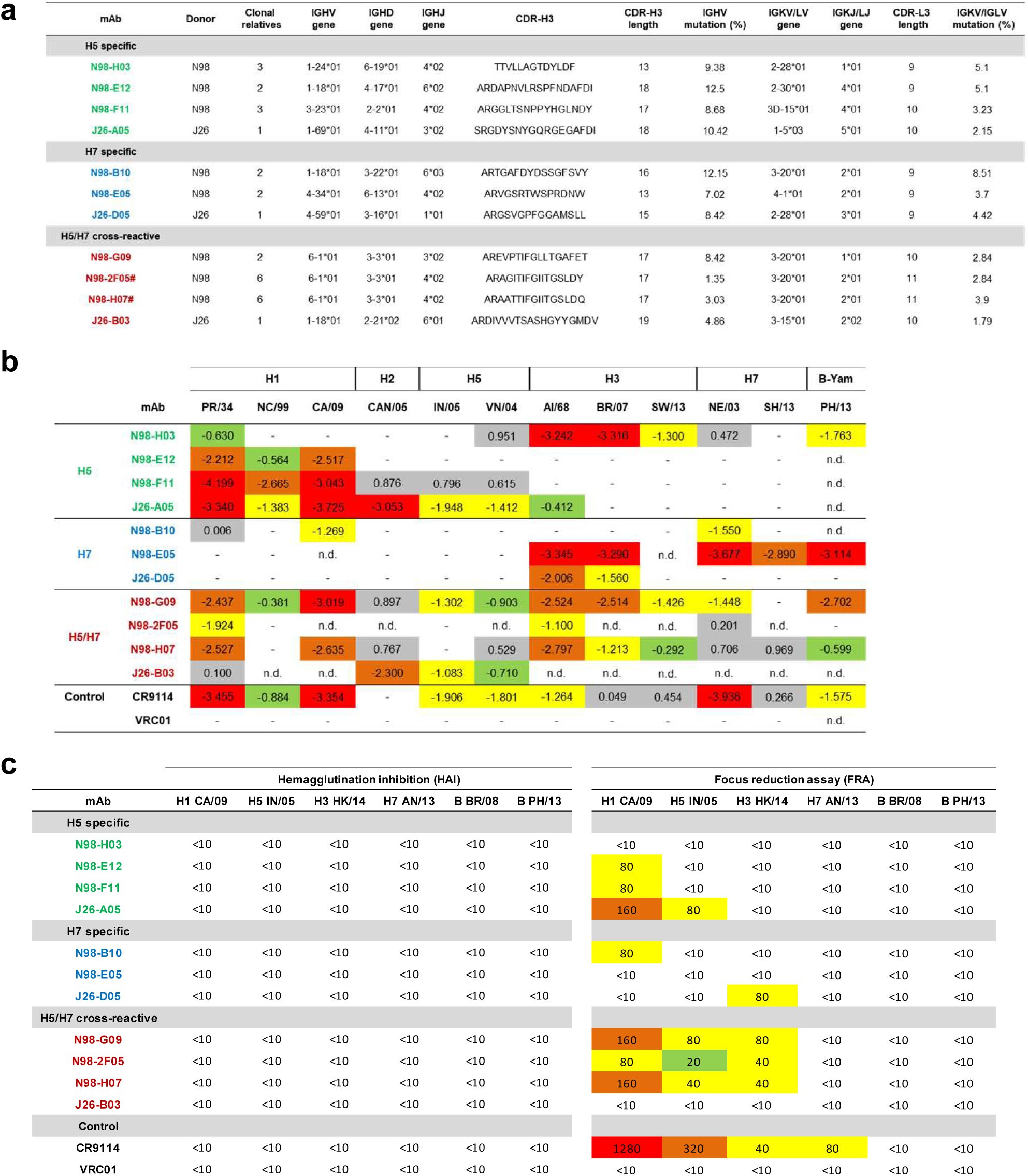
Characteristics of monoclonal antibodies derived from avian HA-specific B cells. (A) Gene usage, CDR-H3 sequence, and mutation loads for 14 monoclonal antibodies derived from sorted H5+, H7+ or H7+H5+ memory B cells from two healthy volunteers. (B) Binding activity of each monoclonal antibody to a panel of recombinant HA proteins derived from influenza A and influenza B strains. Values denote binding log_10_(EC50) determined by ELISA. HA and mAb combinations not tested are indicated by “n.d.” (C) Neutralisation activity determined by hemagglutination inhibition (HAI) or focus reduction assay (FRA) against a panel of influenza A and B viruses. PR/34;A/Puerto Rico/8/1934, NC/99; A/New Caledonia/20/1999, CA/09; A/California/04/2009, HK/68; A/Hong Kong/1/1968, WY/03; A/Wyoming/3/2003, PE/09; A/Perth/16/2009, SW/13; A/Switzerland/9715293/2013, VN/04; A/Vietnam/1203/2004, IN/05; A/Indonesia/5/2005, NE/03; A/Netherlands/219/2003, AN/13; A/Anhui/01/2013, SH/13; A/Shanghai/01/2013

### Responsiveness of memory B cells binding avian HA to vaccination or infection with seasonal influenza viruses

Highly cross-reactive serum antibody responses that bind avian influenza strains are reported to be poorly elicited by inactivated seasonal influenza vaccines^17^. However, the extent to which cross-reactive memory B cells are directly elicited by seasonal vaccines remains unclear. Given the recent spread of avian Clade 2.3.4.4b H5N1 virus, we sought to characterise cross-reactive humoral responses to this H5 clade following administration of the 2017 Southern Hemisphere inactivated quadrivalent influenza vaccine (IIV4).

As expected, serum HAI titres against A/Michi (vaccine component strain) significantly increased 1 month after immunisation (Fig 3A). However, no measurable HAI titres were detected against H5 (A/Fujian-Sanyuan/21099/2017) prior to or after seasonal vaccination. The frequency of class-switched H1+, H5+, and H1+H5+ B cells was assessed in individuals (N=23) using B cell probes derived from H1 (A/Victoria/2570/2019) and H5 (A/Fujian-Sanyuan/21099/2017). H1+ responses significantly increased at 1 month post-immunisation (p=0.0431) although the magnitude of the change in frequency was small (median 1.3-fold relative to baseline) (Fig 3B). No significant changes in H5+ or H1+H5+ responses were observed up to 1 month post-vaccination, with only 3 individuals showing a notable increase in H5-binding cells post-vaccination. The distribution of antibody isotypes and B cell subsets (as defined by CD21 and CD27 expression) of H1 and H5 reactive B cells were broadly similar to the total memory B cell population (Supplementary Fig 2).

**Figure 3:**
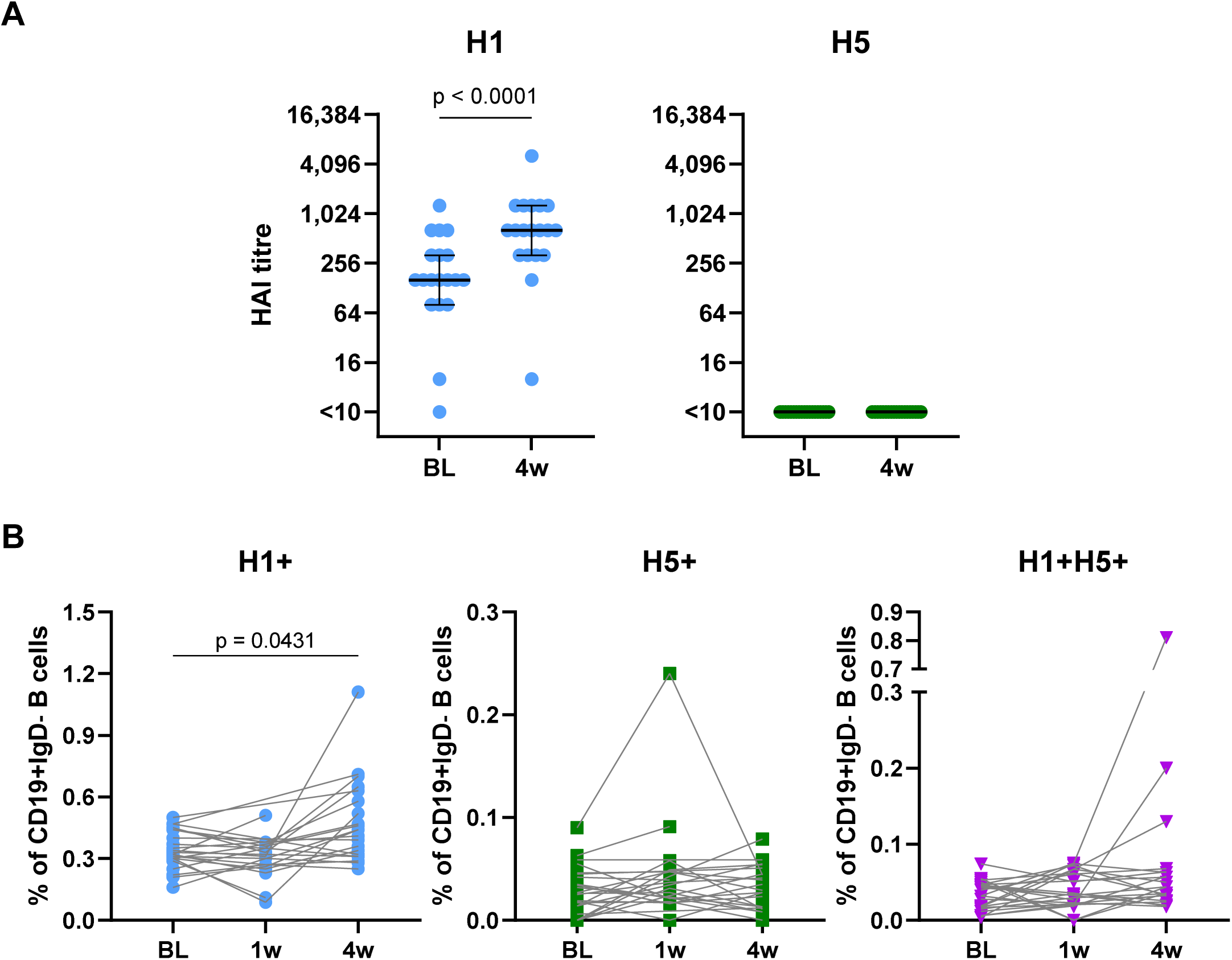
Cross-reactive serum antibodies not induced by seasonal vaccination but sporadic induction of cross-reactive H5 memory B cells. (**A**) Plasma samples from healthy volunteers (N=19) were assayed for HA inhibition (HAI) against the vaccine matched H1N1 strain (H1/Mic) and clade 2.3.4.4b H5N1 strain (H5/Fuj). Baseline (BL) and 4 weeks post immunisation (4w) timepoints were assayed. The lowest plasma dilution assayed was 1:10, with samples not achieving inhibition at this dilution shown as “<10”. The horizontal black line represent the median, and error bars equal the IQR. Significance was determined by Wilcoxon test between BL and 4w. (**B**) Frequencies of H1+ (H1/Vic), H5+ (H5/Fuj) and H1+H5+ memory B cells in healthy volunteers (N=19) at BL, 1- and 4-weeks post vaccination (1w and 4w). Donor responses are each timepoint are linked by lines. Significance was determined by Kruskal-Wallis test with Dunn’s post-test comparing each post-immunisation timepoint to baseline (p values corrected for multiple comparisons).

### CD4+ T cell responses against avian H5 are expanded by seasonal vaccination

CD4+ T cell responses towards H1 and H5 were assessed via ex vivo restimulation and activation-induced marker (AIM) assay. Vaccine-induced expansion of H1-specific CD4+ cells was primarily observed in the cTFH (CD45RA−CXCR5+) compartment (Supplementary Fig 3), with a 4.6-fold increase in median CD184−CD137+ frequency^51^ between baseline and week 1 (p=0.0632), before returning to baseline (Fig 4A). Limited responsiveness was observed within Tmem (CXCR5− and not CCR7+CD45RA+) cells based upon either CD184−CD137+ or CD154 expression. H5-specific CD184−CD137+ Tmem responses were 3.2-fold higher between baseline and 4 weeks post-vaccination (p=0.0419). However, there were no significant changes in cTFH responses following IIV4 immunisation.

**Figure 4:**
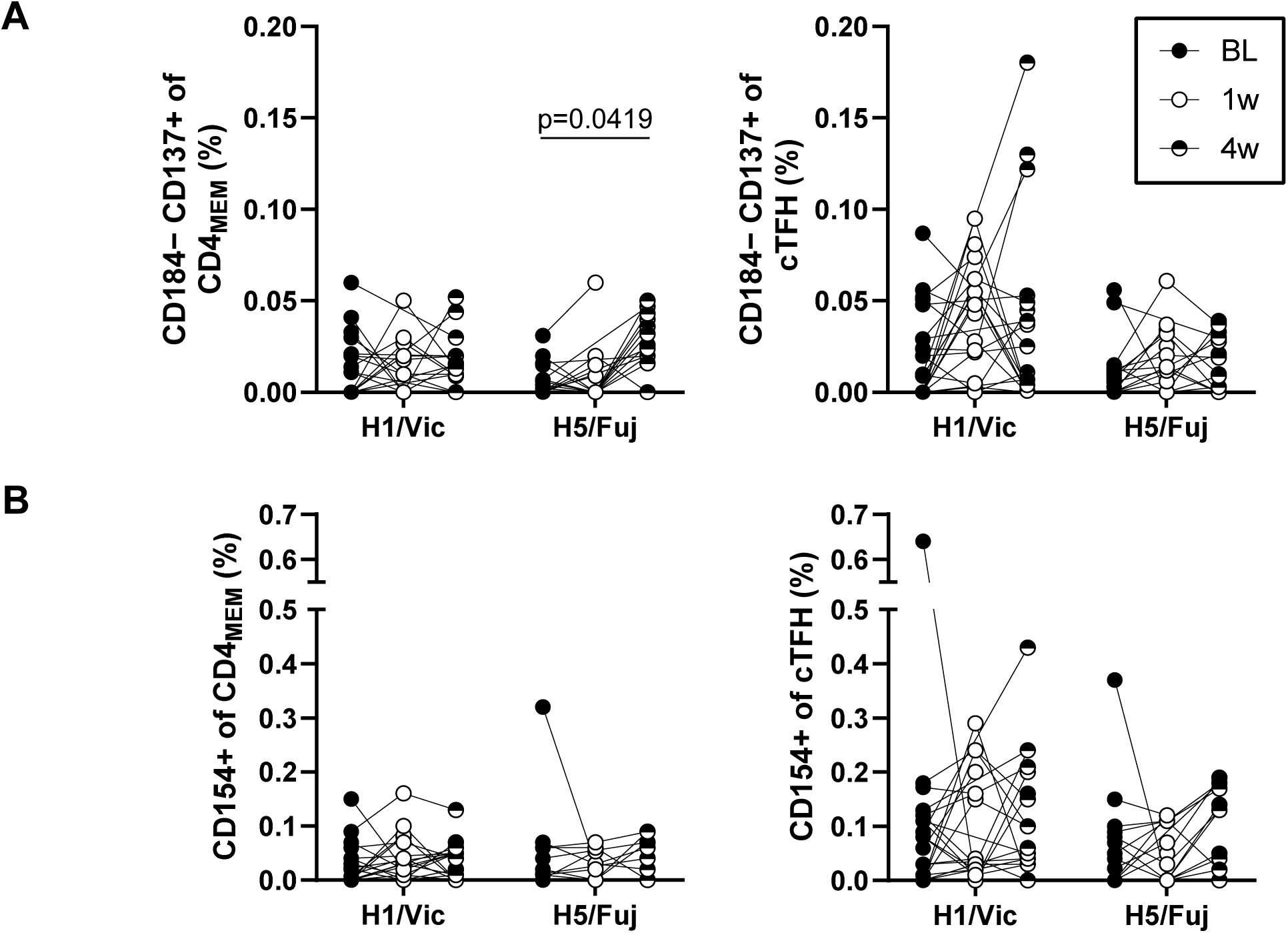
T cell responses against H1 and H5 HA following seasonal vaccination. PBMCs collected from healthy volunteers (N=23) at baseline (BL; black circle) and following seasonal vaccination at 1 week (1w; open circle) and 4 weeks (4w; half-filled circle) post-immunisation were tested via AIM assay against H1/Vic and H5/Fuj HA protein. Antigen-specific CD4+ memory T cells (CD4_MEM_) and circulating T follicular helper cells (cTFH) were identified via (**A**) CD184−CD137+ or (**B**) CD154+ gating. Individual donor responses are plotted and linked by lines between timepoints. Significance was determined by Kruskal-Wallis testing with Dunn’s post-test comparing each post-immunisation timepoint to baseline (p values corrected for multiple comparisons).

## Discussion

The rapid global spread of pathogenic avian strains such as H5 2.3.4.4b has highlighted the omnipresent threat of an influenza pandemic. Highly cross-reactive T cell^27–31^ and B cell^14–16^ populations have been described in influenza-exposed adults and may both a source of background immune protection in the face of a pandemic, or a potential target for expansion via vaccination to broaden immune protection. Here we show that memory B cells with cross-reactive recognition of avian H5 and H7 influenza viruses are detectable in unexposed Australian adults and express immunoglobulins capable of heterosubtypic HA recognition, some with a degree of neutralising activity. As expected and consistent with prior reports, B cells with broadly cross-reactive recognition of influenza A likely to target the HA stem region and are generally drawn from well characterised stereotypic classes (e.g. VH6-1^26,49^, VH1-69^52–54^, VH1-18^26^). At high concentrations, such antibodies have shown an ability to protect against pathogenic infection in pre-clinical challenge settings^17,55–57^ including against H5 2.3.4.4b^58^.

The protective potential against avian influenza offered by routine seasonal vaccination appears relatively limited, based upon low titres of serum antibody recognising HA from avian strains, no baseline HAI activity, and low frequencies of HA-specific B and T cells. While some individuals show evidence of limited cross-reactive immunity to these unencountered HA antigens, confirming previous reports of limited baseline immunity at a population level^59–62^. Upon administration of IIV4, we observed transient boosts in the frequency of H1-specific B and CD4+ T cells as expected. However, this was not recapitulated with regards to H5-specific responses, where expansion of H5 2.3.4.4b responses was limited.

Similarly, our data demonstrate negligible frequencies of H5 2.3.4.4b HA-specific CD4 memory at baseline in a healthy adult cohort, with only minimal augmentation by seasonal vaccination. T cell cross-reactivity between seasonal and avian influenza strains is likely to be greater for conserved internal proteins such as NP or M than for HA, which may constitute a degree of baseline protection in the event of an outbreak. HA-based vaccines (whether split, recombinant protein, or mRNA/LNP), however, rely entirely on T cell help derived from the HA antigen. Our data and others suggest the pool of cross-reactive CD4 T cells established by seasonal influenza exposure is limited, and that pre-priming is required to support the immunogenicity of H5 vaccines^33^. Recently, covalent linkage of H5 HA to seasonal HA proteins successfully augmented H5 antibody responses in human tonsil organoids by “borrowing” existing CD4 memory^36^. Establishment of broad population immunity against pre-pandemic H5 strains may thus require multiple vaccine doses to establish a sufficient pool of T cell help, or rational design of novel vaccines that maximise availability of preexisting CD4 T cell immunity.

Overall, our findings highlight the need for avian influenza-specific vaccine products to bolster immunity in human populations. Vaccines targeting avian influenza strains with pandemic potential have been developed and are immunogenic in humans^7,22,63^. However better pre-pandemic preparedness might necessitate consideration of prophylactic immunisation of human populations prior to an outbreak to expand baseline frequencies of H5- and H7-specific memory T and B lymphocytes, albeit with the understanding that strain matching to the strain that facilitates sustained human-to-human transmission might be imperfect.

## Materials and Methods

### Participant recruitment and sample collection

Study protocols were approved by the University of Melbourne Human Research Ethics Committee (Projects 432/14 and 11395), and all associated procedures were carried out in accordance with the approved guidelines. All participants provided written informed consent in accordance with the Declaration of Helsinki. Participants were not compensated for their participation.

Peripheral blood samples were collected from a cohort of 18 healthy adults as well as a group of 23 healthy individuals who provided samples at baseline, 1 week, and 4 weeks after immunisation with the 2017 Southern Hemisphere inactivated quadrivalent influenza vaccine (IIV4). Whole blood was collected in sodium heparin anticoagulant. Plasma was collected and stored at −80C. Peripheral blood mononuclear cells (PBMC) were collected by Ficoll-Paque separation, washed, and cryopreserved in 10% DMSO/90% fetal calf serum (FCS). PBMC were stored in liquid nitrogen until use.

### HA-specific probes and flow cytometry

The design and purification of fluorescently labelled recombinant HA probes with ablated sialic acid binding activity has been previously described ^47^. HA-specific B cells were identified within cryopreserved PBMC samples by co-staining with relevant combinations of: H7 (A/Shanghai/01/2013), H1 (A/California/04/2009), H1 (A/New Caledonia/20/1999), H3 (A/Hong Kong/1/1968), H3 (A/Victoria/361/2011) H5 (A/Indonesia/05/2005) probes conjugated to streptavidin-PE or -APC (Life Technologies, New York, NY) respectively. B cells were characterised using the following: CD3-QD655, CD14-QD800, CD27-QD605 (Invitrogen), CD19-ECD (Beckman Coulter), IgM-Cy5.5-PerCP, IgG-FITC (BD Pharmingen).

For the seasonal influenza vaccine cohort, B cells were co-stained with H1 (A/Victoria/2570/2019) and H5 (A/Fujian-Sanyuan/21099/2017) probes conjugated to streptavidin-APC or -PE, respectively. The staining panel included IgM BUV395 (G20-127), CD21 BUV737 (B-ly4), IgG BV786 (G18-145), IgD PE-Cy7 (IA6-2) (BD), CD27 BV605 (O323; BioLegend), CD19 ECD along with BV510 dump makers (CD14, M5E2; CD3, OKT3; CD8α, RPA-T8; CD16, 3G8; CD10, HI10a; all from BioLegend) and unconjugated streptavidin-BV510 (BD). For all samples, cell viability was assessed using Aqua Live/Dead amine-reactive dye (Invitrogen). 1–2 million events were collected on an LSR II instrument (BD Immunocytometry Systems) or a FACSymphony A5 SE (BD). Analysis was performed using FlowJo software version 9.5.2 or 10.10 (TreeStar).

### Activation Induced Marker (AIM) assay

Cryopreserved PBMC samples were thawed and rested for 2–4 hr at 37°C in RPMI-1640 supplemented with penicillin/streptomycin/L-glutamate and 10% FCS (Sigma) (RF10). Cells were cultured at 1–2 million cells per well in 200μL in 96-well plates (Corning) and stimulated for 20 hr with 5μg/mL of protein (BSA, A/Victoria/2570/2019 HA or A/Fujian-Sanyuan/21099/2017 HA). Small pools of selected donors were also stimulated with SEB (5μg/mL) as a positive control. Following stimulation, cells were washed and stained with monoclonal antibodies (mAbs) CD183 PE-Dazzle594 (G02H57, BioLegend), CD184 BUV395 (12G5, BD), and CD185 PE-Cy7 (MU5UBEE, ThermoFisher) for 30 min at 37°C. Cells were then washed, stained with Live/Dead Aqua viability dye (ThermoFisher) and incubated with mAbs CD3 BUV805 (SK7), CD20 BV510 (2H7), CD154 PE (TRAP1), CD45RA R718 (HI100) (BD), CD137 BV421 (4B4-1), CD14 BV510 (M5E2), CD4 BV605 (RPA-T4), CD196 BV785 (G034E3), CD134 PerCP-Cy5.5 (ACT35), CD197 Ax647 (G043H7) (BioLegend) for 30min at 4°C. Cells were washed, fixed with 1% formaldehyde and acquired on a BD FACS Symphony A5 SE and analysis was performed using FlowJo Software v10.10 (BD).

### Sequencing, cloning and expression of B cell immunoglobulins

The sequencing and cloning of BCRs from single B cells was performed as previously described^48,64^. Plasmids expressing heavy and light immunoglobulin chains were transfected into Expi293F cells using ExpiFectamine (Invitrogen). Recombinant monoclonal antibodies were purified from culture supernatants using Protein-A or G (Pierce) as per the manufacturer’s instructions.

### Enzyme-linked immunosorbant assay (ELISA)

Antibody binding to HA was tested by ELISA. 96-well Immunosorp plates (Nunc) were coated overnight at 4 °C with 2 μg/mL recombinant HA either expressed in house in Expi293 cells or sourced commercially (Sino Biological). After blocking with 1% fetal calf serum (FCS) in PBS, duplicate wells of monoclonal antibodies (starting at 10 μg/mL, four times serial dilutions) or human sera (1:100, four times serial dilutions) were added and incubated for one hour at room temperature. Plates were washed prior to incubation with 1:20 000 dilution of HRP-conjugated anti-human IgG (KPL) for 1 h at room temperature. Plates were washed and developed using 3,3′,5,5′-Tetramethylbenzidine (TMB) substrate and read at 450 nm. HA-binding activity of monoclonal antibodies was calculated as the antibody concentration giving half-maximal signal (EC50) using a fitted curve (4 parameter log regression). For serum samples, endpoint titres were using a fitted curve (4 parameter log regression) and a cutoff of two times background.

### HAI Assay

HAI was performed according to the WHO Global Influenza Surveillance Network protocols^65^ with the exception that volumes were reduced to 25 µL of ferret sera, 25 µL of antigen (4 HA units), and 25 µL of 1% turkey erythrocytes (0.33% final concentration). Samples were treated with receptor-destroying enzyme (Denka Sieken) at a 1:3 ratio for 18 hours at 37°C, heat inactivated at 56°C for 30min and adsorbed with 5% erythrocytes before testing. The A/Michigan/45/2015 (H1N1) virus was propagated in day-10 embryonated chicken eggs. For H5-ferritin nanoparticles, genes expressing the ectodomain of H5 A/Fujian-Sanyuan/21099/2017 HA were synthesised (IDT-DNA) and cloned into mammalian expression vector allowing the expression of ferritin nanoparticles as described previously^66^. H5-ferritin nanoparticles were expressed using Expi293F cells (ThermoFisher) and purified using HiTrap Anion exchange and CaptoCore chromatography. Purified nanoparticles were diluted to 4 HA units for use in HAI assays.

Assessment of HAI activity of recombinant mAbs was assessed using 1% turkey erythrocytes in a WHO standardised assay. Briefly, mAbs were diluted to 100 μg/mL in PBS prior to incubation with ether treated influenza viruses from strains A/California/07/2009, A/Indonesia/5/2005/PR8-IBDC-RG2, NIBRG-268 A/Anhui/1/2013, A/Hong Kong/4801/2014, B/Phuket/3073/2013 and B/Brisbane/60/2008. HAI titres are reported as the reciprocal of the highest dilution where hemagglutination was completely inhibited.

### FRA Assay

Neutralisation activity of recombinant mAbs against A/California/07/2009, A/Indonesia/5/2005/PR8-IBDC-RG2, NIBRG-268 A/Anhui/1/2013, A/Hong Kong/4801/2014, B/Phuket/3073/2013 and B/Brisbane/60/2008 was examined using focus reduction assays as previously described^67^. The neutralisation titre is expressed as the reciprocal of the highest dilution of a 1 mg/mL mAb stock at which virus infection is inhibited by ≥50%.

### Statistical Analyses

Data is generally presented as median +/− IQR. All statistical analyses were performed using GraphPad Prism v5.0 or v10.4.2 (GraphPad Software Inc.).

## Competing interests

The authors declare no competing interests.

## Authors’ contributions

C.A.G., J.A.J. and A.K.W. designed the study and experiments. C.A.G., M.K., R.E., M.Z.M.Z., Y.L., A.Z., L.S.U.S., D.T., M.A. and A.H. performed experiments. S.J.K. provided unique samples. C.A.G., M.K., J.A.J. and A.K.W. analysed the experimental data. C.A.G., J.A.J. and A.K.W. wrote the manuscript. All authors have read and approved the manuscript.

## Acknowledgements

The authors express gratitude towards the study participants for their provision of samples. We acknowledge the Melbourne Cytometry Platform for provision of flow cytometry services. J.A.J., S.J.K. and A.K.W. were awarded Australian National Health and Medical Research Council Investigator Grants. J.A.J. was the recipient of a Viertel Senior Medical Research Fellowship.

## Notes

### Competing Interest Statement

The authors have declared no competing interest.

