## Supplementary Figures 1-3 for "Memory T and B cells with recognition of avian influenza hemagglutinins are poorly responsive to existing seasonal influenza vaccines"

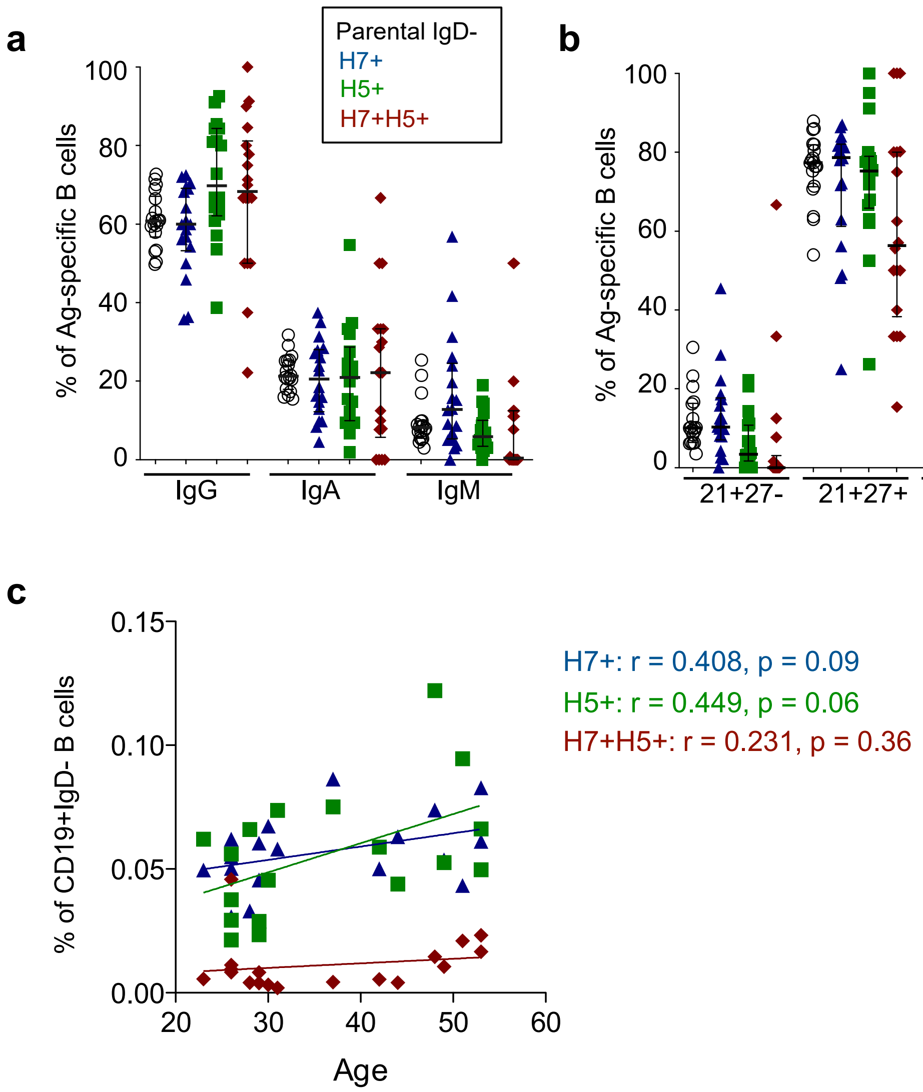
a
Parental IgD-
H7+
H5+
H7+H5+
% of Ag-specific B cells
100
80
60
40
20
0
IgG
IgA
IgM
b
% of Ag-specific B cells
100
80
60
40
20
0
21+27-
21+27+
21-27+
21-27-
c
% of CD19+IgD- B cells
0.15
0.10
0.05
0.00
20
30
40
50
60
Age
H7+: r = 0.408, p = 0.09
H5+: r = 0.449, p = 0.06
H7+H5+: r = 0.231, p = 0.36

**Supplementary Figure 1. Subclass and phenotype of H7 and H5 HA-specific memory B cells.**

(**A**) Distribution of IgG, IgA and IgM expression on H7+ (blue), H5+ (green), H7+H5+ (red) and the parental memory B cell population (open circles) in healthy volunteers (N=18). (**B**) Distribution of B cell subsets delineated by CD27 and CD21 staining: CD27-CD21+ naïve, CD27+CD21+ resting memory, CD27+CD21- activated memory and CD27-CD21- tissue-like populations. Lines indicate median and IQR. **(C)** Correlation between participant age and HA-specific memory B cell frequencies (N=18). Statistics assessed by spearman correlation.

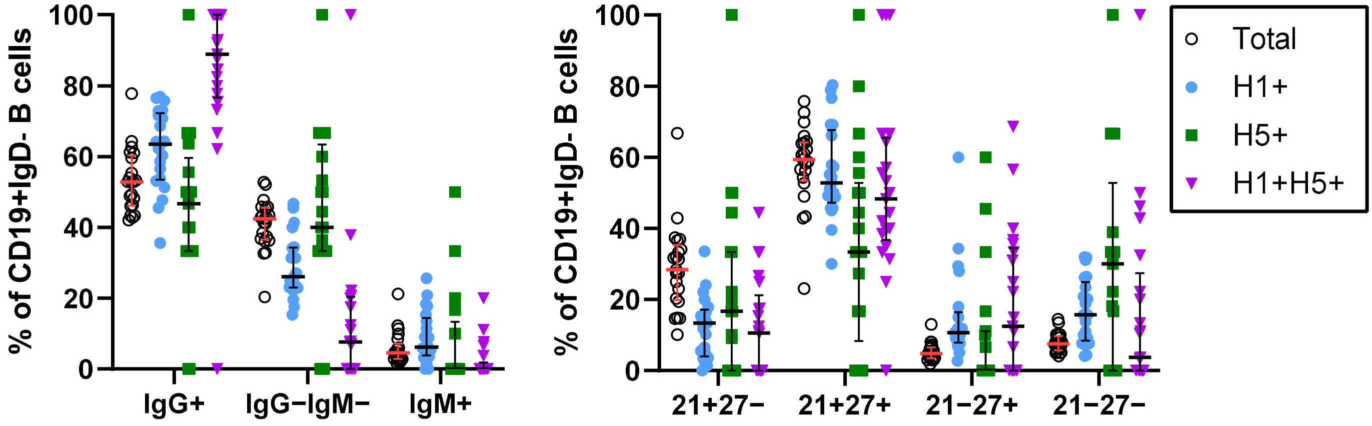

% of CD19+IgD- B cells
100
80
60
40
20
0
IgG+
IgG−IgM−
IgM+
% of CD19+IgD- B cells
21+27−
21+27+
21−27+
21−27−
Total
H1+
H5+
H1+H5+

**Supplementary Figure 2. Subclass and phenotype of H1 and H5 HA-specific memory B cells.**

(**A**) Distribution of IgG, IgG−IgM− and IgM expression on H1+ (light blue), H5+ (green), H1+H5+ (purple) and the parental memory B cell population (total IgD−, open circles) in healthy volunteers (N=23). (**B**) Distribution of B cell subsets delineated by CD27 and CD21 staining: CD27−CD21+ naïve, CD27+CD21+ resting memory, CD27+CD21− activated memory and CD27−CD21− tissue-like populations. Horizontal black lines indicate the median and error bars equal the IQR (lines coloured red for “total IgD−” for clarity).

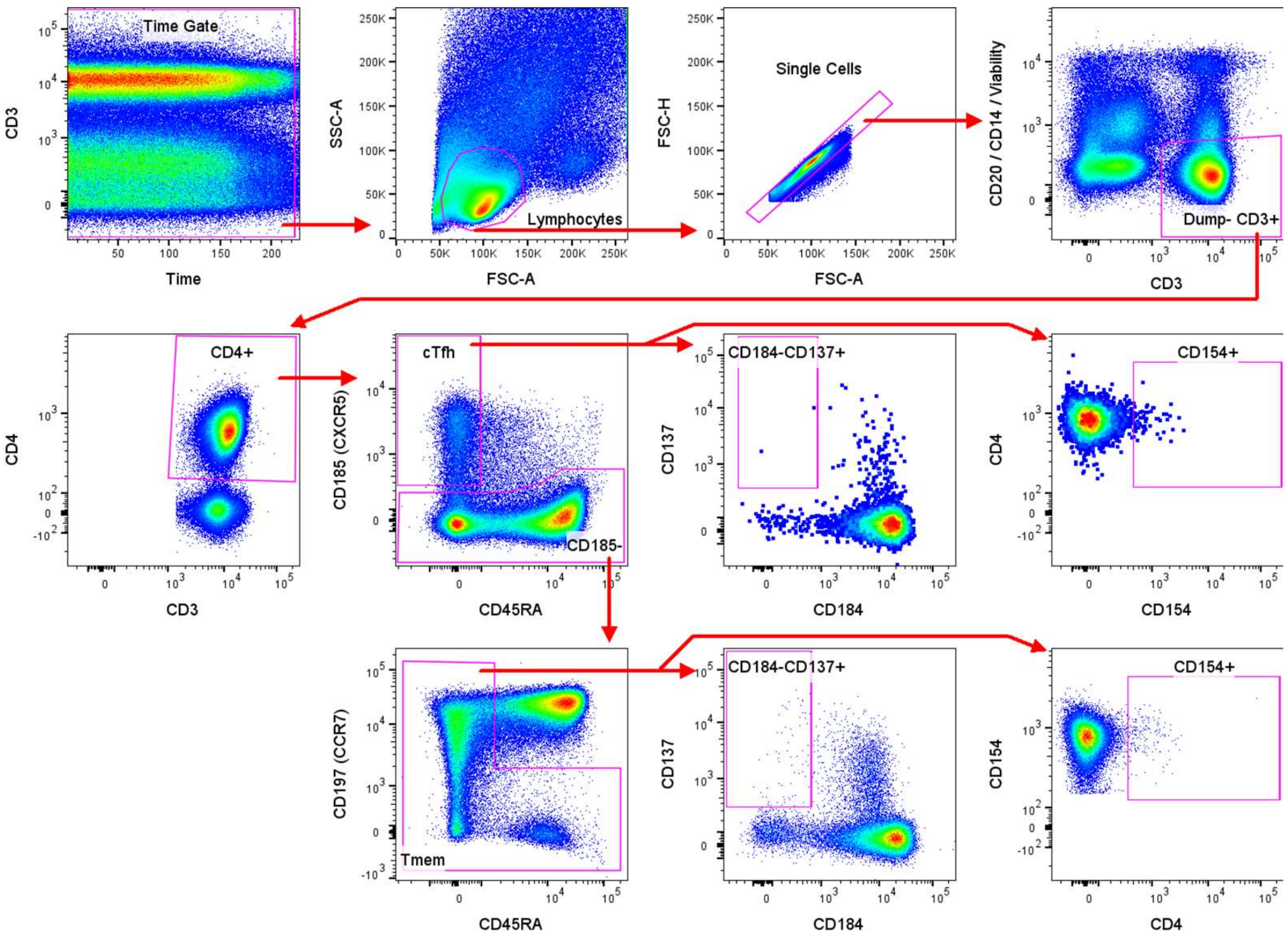
Time Gate
CD3
Time
SSC-A
FSC-A
Lymphocytes
FSC-H
Single Cells
CD20 / CD14 / Viability
Dump- CD3+
CD4+
CD4
cTfh
CD185 (CXCR5)
CD185-
CD45RA
CD184-CD137+
CD137
CD184
CD154+
CD154
CD197 (CCR7)
Tmem

**Supplementary Figure 3. Flow cytometry gating for AIM assay.**

Staining of cryopreserved PBMCs to identify circulating T follicular helper (cTFH) cells (CXCR5+) and memory T (Tmem) cells (CXCR5− and not CCR7+CD45RA+). For each T cell subset, antigen specific cells were identified by CD137+CD184− or CD154+ staining.
